# Deep learning–based automated prediction of mouse seminiferous tubule stage by using bright-field microscopy

**DOI:** 10.1101/2024.08.07.606973

**Authors:** Yuta Tokuoka, Tsutomu Endo, Takashi Morikura, Yuki Hiradate, Masahito Ikawa, Akira Funahashi

**Author notes:** These authors contributed equally to this work (Yuta Tokuoka, Tsutomu Endo). Corresponding authors: M.I., A.F.

## Abstract

Infertility is a global issue, with approximately 50% of cases attributed to defective spermatogenesis. For studies into spermatogenesis and spermatogenic dysfunction, evaluating the seminiferous tubule stage is essential. However, the current method of evaluation involves labor-intensive and time-consuming manual tasks such as staining, observation, and image analysis. Lack of reproducibility is also a problem owing to the subjective nature of visual evaluation by experts. In this study, we propose a deep learning–based method for automatically and objectively evaluating the seminiferous tubule stage. Our approach automatically predicts which of 12 seminiferous tubule stages is represented in bright-field microscopic images of mouse seminiferous tubules stained by hematoxylin-PAS. For training and validation of our model, we created a dataset of 1229 tissue images, each labeled with one of 12 distinct seminiferous tubule stages. The maximum prediction accuracy was 79.58% which rose to 98.33% with allowance for a prediction error of ±1 stage. Remarkably, although the model was not explicitly trained on the patterns of transition between stages, it inferred characteristic structural patterns involved in the process of spermatogenesis. This method not only advances our understanding of spermatogenesis but also holds promise for improving the automated diagnosis of infertility.

## Introduction

Infertility is a worldwide social problem, affecting an estimated 8%–12% of couples of reproductive age [1]. Approximately 50% of cases are caused by male factors, the majority of which are recognized as defective spermatogenesis. Although some of the major regulatory genes and environmental factors involved in defective spermatogenesis have been identified[2], the mechanisms underlying the nature of spermatogenesis remain unclear.

Researchers have attempted to gain insights into spermatogenesis through histologic approaches involving microscopic observations of the cellular structure of tissues. A major approach divides the mouse seminiferous epithelial cycle into 12 distinct seminiferous tubule stages serving as landmarks[3]. Identifying these stages enables researchers to determine which differentiation stage of spermatogenesis the germ cells are in. This makes it possible to identify when abnormalities in spermatogenesis occur and to analyze mRNA and protein expression and localization[4]. In general, methods of identifying stages have used immunofluorescence staining to label specific histologic markers or have relied on microscopic observation of sections of seminiferous tubules. However, these methods require a lot of manual work, including staining, observation, and image processing, which are labor-intensive and time-consuming. Lack of reproducibility is also another problem, because the identification of stage can differ between experts.

In recent years, there have been reports of research using deep learning to automate and standardize the analysis of tissue images to overcome problems of high human labor and time costs and low reproducibility[5, 6, 7]. Among such studies are those using deep learning to predict the seminiferous tubule stage. Xu et al. reported their system of Computerized Spermatogenesis Staging (CSS) that uses deep learning to predict the seminiferous tubule stage in bright-field microscopic images of hematoxylin and eosin (H&E)-stained seminiferous tubules[8]. However, the CSS system simplifies the 12 seminiferous tubule stages into three: early stage (Stages I–V), intermediate stage (Stages VI-VIII), and late stage (Stages IX-XII). Yang et al. developed their Software for Analysis of Testis Images with Neural Networks (SATINN), which can perform fine-grained tubule stage prediction from immunofluorescence-stained images by using deep learning[9]. However, although SATINN is able to classify seminiferous tubules into 12 stages, it requires immunofluorescence-stained images. Not only does this limitation impose human labor and time costs, but also reproducibility can be degraded owing to photobleaching of the immunofluorescence-stained images. In addition, it precludes any subsequent downstream analysis such as H&E image analysis.

In this study, we developed a deep learning method that can automatically and objectively predict the seminiferous tubule stage (from among 12 stages) that is represented in a bright-field image of seminiferous tubules. For training and evaluation of the learning model, we created a dataset by assigning one of 12 stages to each of 1229 seminiferous tubule tissue images captured by bright-field microscopy. We used four neural network architectures as prediction models: ResNet-50[10], ResNeXt-50[11], WideResNet-50[12], and MobileNet-v3[13]. These four models were evaluated by four-fold cross-validation to assess the accuracy of their seminiferous tubule stage prediction. The trained models obtained by cross-validation were then validated by using test data. Our best-performing model achieved a maximum stage prediction accuracy of 79.58%, and when the prediction error was allowed to be ±1 stage, the prediction accuracy was 98.33%. Furthermore, the trained model was found to predict the stages via unique cell shapes in seminiferous tubules. This is consistent with the features that experts look for when classifying seminiferous tubule stages, suggesting that the trained model predicts the stages on a basis similar to that of a skilled expert. More interestingly, even though the temporal sequence of stage transitions during spermatogenesis was not explicitly given, our model learned the transition patterns of the characteristic structures involved in spermatogenesis. Our method not only contributes to the understanding of spermatogenesis but also may also be useful in the diagnosis of male infertility.

## Methods

### Mice

All animal experiments were approved by the Animal Care and Use Committee of the Research Institute for Microbial Diseases, Osaka University (#Biken-AP-H30-01). We complied with all relevant ethical regulations for animal use. Wild-type C57BL/6J mice were purchased from Charles River Laboratories Japan, Inc. (Yokohama, Japan) and 6N mice were purchased from CLEA Japan, Inc. (Tokyo, Japan).

### Identification of the stages of the seminiferous cycle

Testes were collected from 2-month-old mice, fixed in Bouin ‘s solution, embedded in paraffin, sectioned, and stained with hematoxylin and periodic acid–Schiff (He-PAS). All sections were observed under a light microscope. Seminiferous tubule stages in the He-PAS–stained sections were determined according to morphological criteria[3, 14], which were the same as those reported previously[15, 16, 17]. In brief, the 12 stages were identified primarily on the basis of the first 12 steps of spermatid development. Germ cell types were identified by their location, nuclear size, and chromatin pattern[14].

### Seminiferous tubules dataset

Tissue images of He-PAS–stained seminiferous tubules were captured with a bright-field microscope to determine the shapes of the nuclei and the apical part. We used an Olympus BX53 bright-field microscope, an Olympus DP80 camera, and an Olympus cellSens Dimension 1.18 for image acquisition. The resolution of the microscopic images was 170.635 *nm*/pixel. Next, we cropped the tissue images to 2040 × 1536 pixels so that the seminiferous tubules appeared in the center, and obtained 1229 RGB images. We then assigned a stage to each seminiferous tubule in the acquired images according to the criteria[3, 14, 15, 16, 17] (Table 1, Supplementary Figure S1).

**Table 1:** Details of the seminiferous tubule stages in the dataset.

| Stage | I | II | III | IV | V | VI | VII | VIII | IX | X | XI | XII | total |
| --- | --- | --- | --- | --- | --- | --- | --- | --- | --- | --- | --- | --- | --- |
| Number of images | 101 | 104 | 101 | 102 | 104 | 105 | 102 | 101 | 103 | 103 | 101 | 102 | 1229 |

We randomly selected 20% of the dataset as test data in advance and used the remaining data as training data. The training data were then randomly divided into four parts for cross-validation. The splitting of datasets was performed in such a way that the proportions of each stage were uniform.

### Details of the neural networks

For precise seminiferous tubule stage prediction we used convolutional neural networks (CNNs), which are neural networks specialized for image-related tasks. We employed ResNet-50[10], ResNeXt-50[11], WideResNet-50[12], and MobileNet-v3[13], which are CNN architectures that can perform image classification tasks with high accuracy (Figure 1). ResNet has a residual module that allows for effective training of deep neural networks.

**Figure 1:**
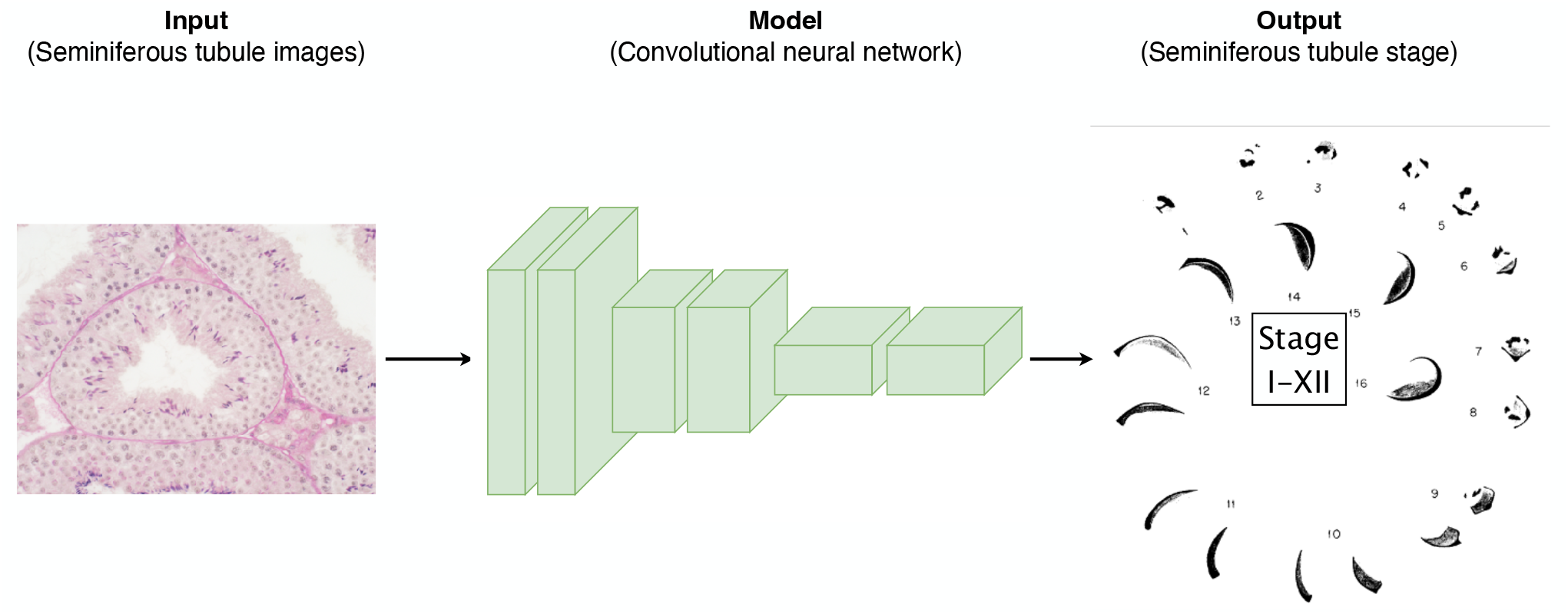
Conceptual diagram of seminiferous tubule stage prediction (adapted from [3]) The diagram represents a convolutional neural network (CNN)-based model whose input is a tissue image with a seminiferous tubule at the center and whose output is a fine-grained classification of the tubule stage (12 levels). ResNet-50, ResNeXt-50, WideResNet-50, and MobileNet-v3 were used for the CNN models.

ResNeXt can learn various feature representations by introducing a mechanism to adjust the depth and width of the network on the basis of ResNet. WideResNet is another ResNet-based model that achieves high accuracy by increasing the network ‘s width without increasing its depth. MobileNet is a model whose lightness (low degrees of freedom) and image classification performance are optimized by Neural Architecture Search[18]. These models have been widely used for image classification tasks.

As the classification task is to predict one of 12 seminiferous tubule stages, the output layer of each CNN model consists of a 12-dimensional fully connected layer. Softmax cross-entropy was used as the objective function.

### Training procedure

For seminiferous tubule stage prediction, the deep learning models were trained on images labeled with each of the 12 different stages. In this case, we trained the models so that all labels (12 stages) were always included in the mini-batch.

The hyperparameters used for training were the same for all models (Supplementary Table S1). We trained for 5000 epochs and used Adam[19] (learning rate = 0.001) as the optimizer. The mini-batch size was set to 12 and the weight decay was set to 0.0005. Min-max normalization was used to preprocess the image intensity values. In addition, random flips, rotations, and crops were conducted to augment the training data. For random flips, the input images were randomly flipped vertically or horizontally at each epoch. For random rotations, the input images were randomly rotated by 90 degrees at each epoch. For random crops, the patch size was set to 896 × 896 pixels. We used an NVIDIA Tesla V100 GPU (operating frequency, 1370 MHz; single precision floating point performance, 14.0 TFLOPS) for training and inference.

### Evaluation metrics and methods

To evaluate the accuracy of the tubule stage prediction, we used the evaluation metrics Balanced Accuracy and F-measure. Balanced Accuracy and F-measure are expressed as equations (1) and (2):

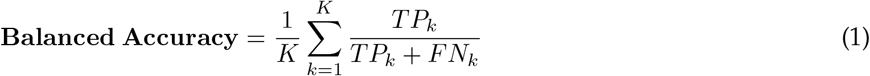

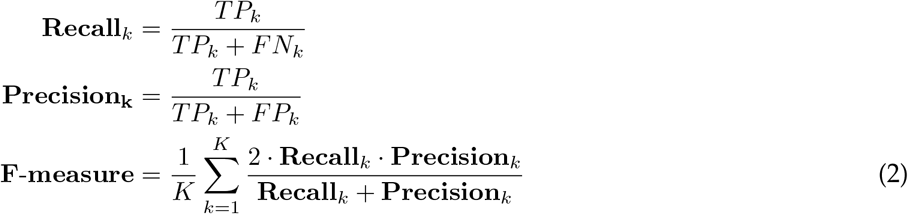

where *K* represents the number of classes, *TP*_*k*_ represents the number of true positives in class *k, FN*_*k*_ represents the number of false negatives in class *k*, and *FP*_*k*_ represents the number of false positives in class *k*. Balanced Accuracy is a metric that evaluates the correct response rate independent of class bias by normalizing the correct response rate of a prediction by the number of samples in each class. F-measure is a metric that can evaluate both false positives and false negatives by harmonically averaging the Recall, which evaluates the absence of false negatives in a prediction, and the Precision, which evaluates the absence of false positives.

We evaluated each of the trained models by four-fold cross-validation. In addition, we selected the model at the epoch with the highest accuracy (Balanced Accuracy) from the validation data in the four-fold cross-validation as the convergent model, and we evaluated this convergent model with the test data.

### Model feature analysis

To analyze the features important to the trained models, we used Grad-CAM (Gradient-weighted Class Activation Mapping)[20], which creates a heat map for visualizing and understanding which features the model is focusing its attention on when making predictions in an image classification task. The higher the value in the heat map, the more important is that feature in contributing to the prediction. In this study, we applied Grad-CAM to the trained ResNet-50 model.

To analyze the features of the trained model, we extracted the features computed in the connection from the convolutional layer to the fully connected layer of the trained model. Features extracted from the ResNet-50 model, which was trained on the seminiferous tubule image dataset constructed in this study, were compared with features from ImageNet[21], which was trained on a general natural image dataset. We visualized the distribution of the features by dimensionality reduction using t-SNE (t-distributed stochastic neighbor embedding)[22].

## Results

### Cross-validation of seminiferous tubule stage prediction models

To validate the performance of our models, we first evaluated their learning curves by four-fold cross-validation using the training dataset (Figure 2). For all four validated models, we found that cross-entropy loss and Balanced Accuracy in the training dataset reached a plateau at approximately 1000 epochs. In the validation dataset, on the other hand, we observed that the loss and Balanced Accuracy had a variance larger than in the training dataset and that accuracy was highest after 1000 epochs.

**Figure 2:**
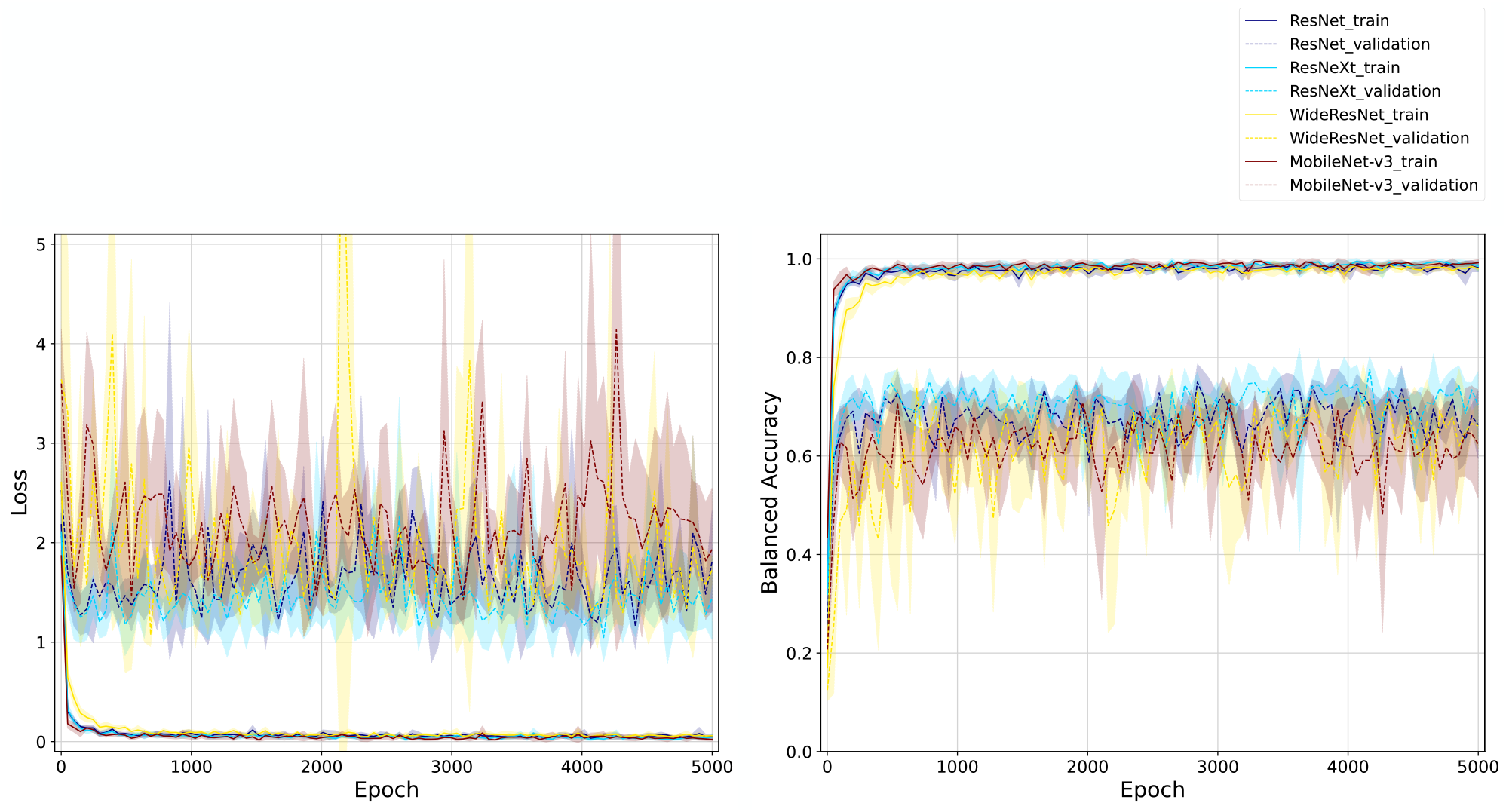
Learning curves for seminiferous tubule stage prediction models. The graphs plot the evolution of loss (left panel) and balanced accuracy (right panel) per epoch in the training of the seminiferous tubule stage prediction models. Each series represents the learning curves for four different models (ResNet-50, ResNeXt-50, WideResNet-50, and MobileNet-v3) and their training (solid line) and validation (dashed line) subsets. The shaded area in each series represents the standard deviation.

We compared the performance of these models with the highest Balanced Accuracy on the validation dataset and found that ResNeXt-50 had the highest accuracy (Balanced Accuracy = 0.8174) (Supplementary Table S2). The other models also achieved a Balanced Accuracy of 0.78 or better. The low standard deviation of the Balanced Accuracy for cross-validation for each model showed that there was no label imbalance or image quality bias in the dataset. These results indicated that all four validated models could learn to predict the stages with high accuracy.

### Test performance evaluation of the seminiferous tubule stage prediction model

We evaluated the performance of the seminiferous tubule stage predictions on the test data by using each of the models with the highest Balanced Accuracy in the validation data set in the four-fold cross-validation (Table 2). All models achieved a Balanced Accuracy and F-measure of 0.7 or higher. Among them, WideResNet-50 had the highest prediction accuracy (Balanced Accuracy = 0.7958, F-measure = 0.7929).

**Table 2:** Accuracy of seminiferous tubule stage prediction of each model using the test dataset. Bold type indicates the maximum value at each metric.

| Model | Balanced Accuracy | F-measure |
| --- | --- | --- |
| ResNet-50 | 0.7833 | 0.7834 |
| ResNeXt-50 | 0.7792 | 0.7781 |
| WideResNet-50 | **0.7958** | **0.7929** |
| MobileNet-v3 | 0.7250 | 0.7083 |

These performances were evaluated by confusion matrix (Figure 3). For all models, most of the errors between the predicted and correct stages were within ±1 (for example, when the correct answer is ‘2’, the ±1 range is ‘1’ to ‘3’). On the basis of these confusion matrices, we calculated and compared the Balanced Accuracy for each model when the error tolerance was set to ±1 and ±2. All models achieved a Balanced Accuracy of 0.96 or better with an acceptable margin of error (Table 3). Among these models, ResNet-50 achieved a maximum accuracy of 98.33% when allowing for ±1 stage of prediction error, and a maximum accuracy of 100% when allowing for ±2 stages of prediction error.

**Table 3:** Balanced Accuracy of each model in predicting the seminiferous tubule stage using the test dataset when errors were allowed. Bold type indicates the maximum value at each allowable error.

| Model | Stage $\pm 0$ | Stage $\pm 1$ | Stage $\pm 2$ |
| --- | --- | --- | --- |
| ResNet-50 | 0.7833 | **0.9833** | **1.0000** |
| ResNeXt-50 | 0.7792 | 0.9625 | 0.9917 |
| WideResNet-50 | **0.7958** | **0.9833** | 0.9958 |
| MobileNet-v3 | 0.7250 | 0.9708 | 0.9917 |

**Figure 3:**
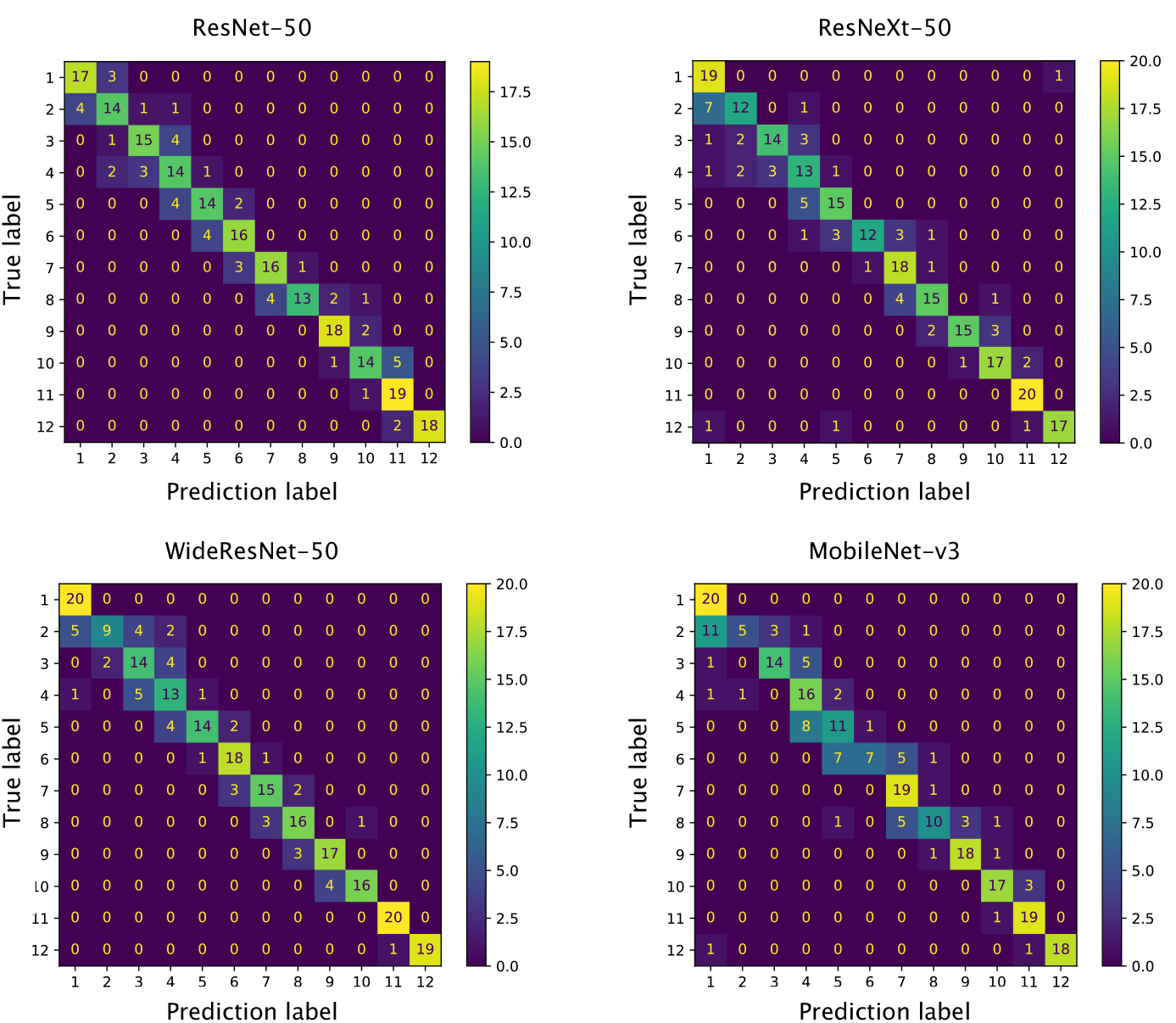
Confusion matrix of seminiferous tubule stage prediction models. The stage prediction results for each test subset are represented by a confusion matrix according to the seminiferous tubule stage prediction model. Confusion matrixes are shown for the predictions made by ResNet-50, ResNeXt-50, WideResNet-50, and MobileNet-v3, respectively. The vertical axis represents the correct stage and the horizontal axis represents the predicted stage.

### Feature analysis of model as the basis for predicting stages

We performed a feature analysis of the trained ResNet-50 model to see which features of the seminiferous tubule stage prediction model were used as the basis for the prediction. Visualization by Grad-CAM[20] showed that the seminiferous tubule features in the images that successfully predicted the stage were circular spermatids, elongated spermatids, and diplotene and meiotically dividing spermatocytes that circularly surround the lumen of the seminiferous tubule (Figure 4, Supplementary Figure S2). Interestingly, the main features that were focused on in the most of seminiferous tubules (stages I-XI) were circular spermatids and elongated spermatids; no attention was paid to spermatogonia or (preleptotene, leptotene, and early zygotene) spermatocytes located near the basement membrane of the seminiferous tubules (Figure 4A–C). On the other hand, in stage XII tubules, the features of interest were elongating spermatids and spermatocytes in the diplotene and late zygotene stages away from the basement membrane of the seminiferous tubules, again with no attention paid to spermatogonia (Figure 4D). These results suggest that the model correctly learned the progression of spermatids into the lumen as they differentiated.

**Figure 4:**
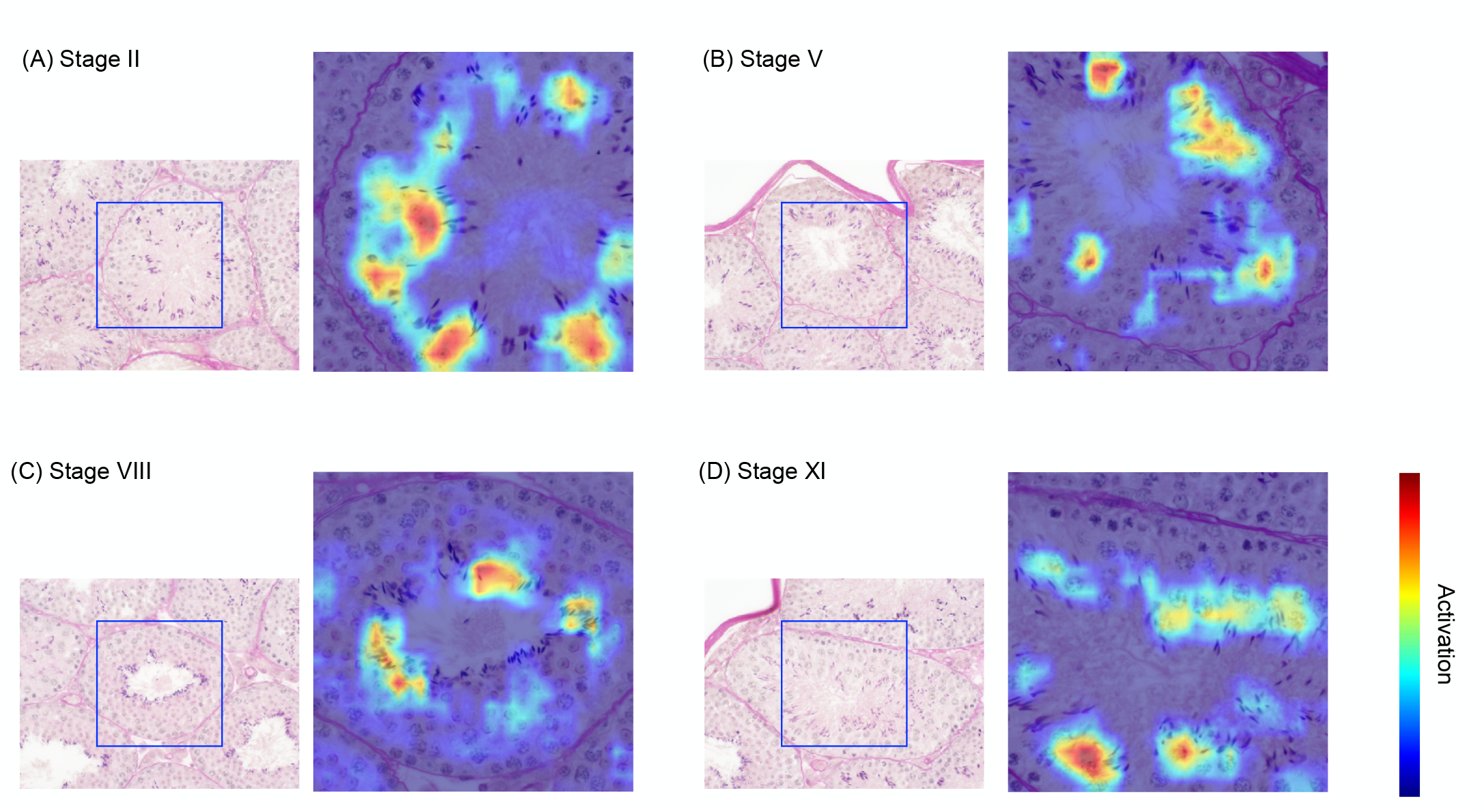
Image features forming the basis for seminiferous tubule stage prediction. Features of the seminiferous tubule images that were used as the basis for stage prediction by ResNet-50 model trained on the seminiferous tubule dataset, visualized by Grad-CAM. All of these results represent the image features that were used as the basis for correct stage prediction. Higher activation in the heat map (indicated by warmer colors) indicates a stronger contribution to the prediction. (A) Features used to predict Stage II. (B) Features used to predict Stage V. (C) Features used to predict Stage VIII. (D) Features used to predict Stage XI.

We also visualized the image features in cases where the seminiferous tubule stage failed to be correctly predicted (Supplementary Figure S3). Three patterns were found in these cases. The first pattern was that the model focused on image features other than the target (centered) seminiferous tubules (Supplementary Figure S3A,B, arrowheads). The second pattern was that the model focused on the lumen of the seminiferous tubules where no spermatogonia, spermatocytes, or spermatids were present (Supplementary Figure S3C,D, arrowheads). The third pattern was that the model focused on only some of the specific cells surrounding the lumen (Supplementary Figure S3E,F). If too many features other than those relevant to the differentiation process appear in the image, the model may output prediction errors.

### Relationships between features and stages in the prediction of seminiferous tubule stage

To gain insight into how the seminiferous tubule stage prediction models learned the important features of seminiferous tubules, we analyzed the relationships between features and stages in the trained ResNet-50 model. By comparing the ResNet-50 model that had learned the features of seminiferous tubule images against the ResNet-50 model trained on general images (ImageNet[21]), we were able to identify patterns in the image features that the model learned (Figure 5). To identify patterns, we used dimensionality reduction by t-SNE[22] to visualize the distribution of the features obtained from the input of seminiferous tubule images at each stage. In the trained ResNet-50 model, the image features formed clusters for each stage (Figure 5A), whereas the results from the model trained on the ImageNet data did not form clusters for each stage (Figure 5B). This indicates that the ImageNet model could not accurately extract the features of the seminiferous tubule stages.

**Figure 5:**
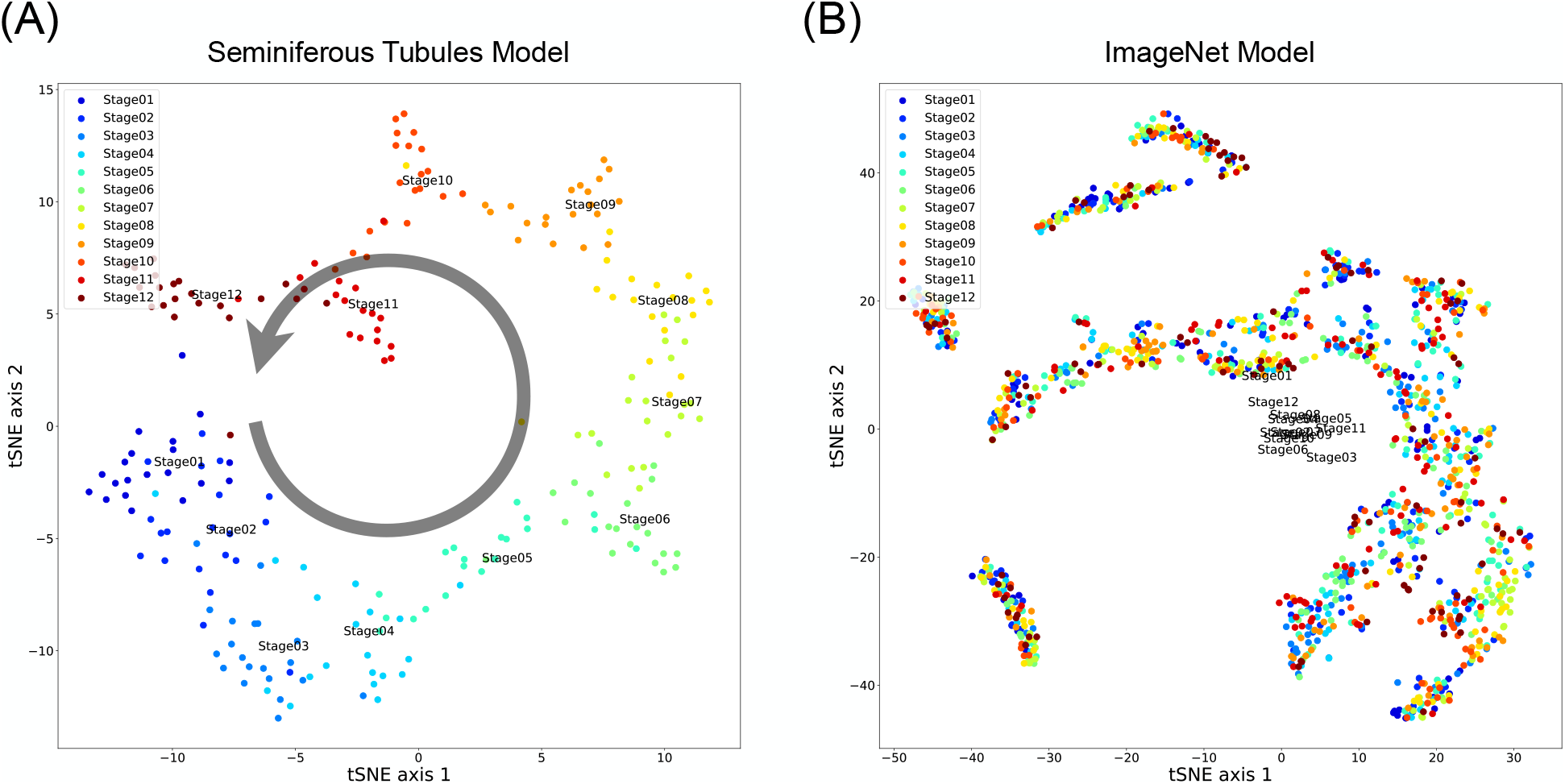
Visualization of feature representations within the seminiferous tubule stage prediction model. Seminiferous tubule image features extracted from ResNet-50 model trained on the seminiferous tubule dataset and ImageNet data, visualized by dimensionality reduction by t-SNE. Each point plotted represents the feature representation of each inputted seminiferous tubule image, and the labels represent the seminiferous tubule stage associated with each image. (A) Visualization of the feature representations used when predicting the seminiferous tubule stage by the model trained on the seminiferous tubules dataset. Small clusters are formed for each seminiferous tubule stage. The arrow indicates the continuous cycling of the stage order. (B) Visualization of the feature representation used when predicting the seminiferous tubule stage by the model trained with ImageNet data. Unlike in (A), clusters are not formed for any of the seminiferous tubule stages.

Focusing on the relationships between features and stages in the model trained on the seminiferous tubule dataset (Figure 5A), it can be observed that a circular structure was formed in which the features of adjacent stages were spaced almost equidistant from each other. This is consistent with the cyclic nature of the seminiferous tubule stages (Stage XII and Stage I are continuous). These results suggest that the model was able to learn that the pattern of seminiferous tubule stages is continuous and cyclical, even though this pattern was not explicitly given in the learning process.

## Discussions

In this study, we developed a CNN-based model capable of automatically and uniformly predicting each of the 12 seminiferous tubule stages from tissue images captured by a bright-field microscope. The model achieved an accuracy of 79.58% in predicting the stage of seminiferous tubules on test data, and when a margin of error of ±1 stage was allowed, the model was able to predict the stage with an accuracy of 98.33%. Feature analysis of the prediction model suggested that the model was able to learn the pattern of continuous change in features that occurs in seminiferous tubules.

Although the seminiferous tubule stage in mice is divided into 12 stages, a single seminiferous tubule may contain a mixture of features from different stages, such that these features may confound an expert ‘s judgment and thus lead to poor reproducibility of stage determination. This is one reason why it is not easy to train machine learning models on the 12-stage classification, discrimination of which can be difficult even for experts. Earlier studies attempted to solve this problem of mixed stage characteristics by restricting the number of stages. For example, the CSS system[8] redefines the 12 stages into 3 stages–early stage (Stages I–V), middle stage (Stages VI-VIII), and late stage (Stages IX-XII)-with stage prediction accuracies reported to be, respectively, 93%, 94%, and 89%. On the other hand, we trained the WideResNet-50 model to perform a fine-grained classification task with 12 stages in an appropriate way and found that it was possible to classify stages with an accuracy of approximately 80%. With ResNet-50 model, we also achieved 98.33% accuracy by allowing a prediction error of ±1 stage and 100% accuracy by allowing a prediction error of ±2 stages. Our method achieved high classification accuracy while remaining close to the conventional method of stage classification.

Another study reported that SATINN, which can predict the stage of seminiferous tubules from immunofluorescencestained images, achieved an accuracy of 94.9% by allowing ±1 stage of prediction error [9]. Therefore, our method is superior to SATINN in terms of accuracy. Furthermore, our method requires only bright-field microscopy, which is superior in that it does not require immunofluorescence staining and its associated human labor and time costs. In addition, unlike SATINN, our method is not prone to the effects of photobleaching or degradation in reproducibility that is associated with immunofluorescence staining. In addition, our method allows subsequent downstream analysis because the seminiferous tubule section can be reused due to the high preservability, unlike the immunofluorescence-stained sections.

Analysis of seminiferous tubule images by using Grad-CAM to identify the features that provide the basis for predicting seminiferous tubule stage revealed that the prediction model mainly focuses on circular spermatids, elongated spermatids, and diplotene and meiotically dividing spermatocytes surrounding the lumen of the seminiferous tubules (Figure 4, Supplementary Figure S2). On the other hand, the model did not focus on features from spermatogonia to spermatocytes in the early zygotene stage, which are important stage-determining characteristics. If we were to train the model with some constraints to force it to learn the features of these spermatogonia and spermatocytes, we could further improve its performance. In addition, we found that the image features that the model focused on in the failed predictions fell into three patterns (Supplementary Figure S3), with the common point among these three patterns being the failure to focus on cells in the seminiferous tubules. From these results, it is clear that paying attention to the cells in the seminiferous tubules in a circular fashion is a prerequisite for accurate prediction of the tubule stage. When experts classify the stages of seminiferous tubules, they identify the spermatogonia, spermatocytes, and spermatids in seminiferous tubules by checking the cell shapes[3, 14, 15]. As the differentiation of spermatids in seminiferous tubules is continuous and coordinated, it makes sense that focusing on only some specific cells that are slightly differentiated or have slightly delayed differentiation could cause a prediction stage error of ±1. These results suggest that our model for predicting the stage of the seminiferous tubules is highly accurate because it focuses on image features that are consistent with the features that experts refer to as the basis for classification.

Analysis of the relationship between features and stages in seminiferous tubule stage prediction revealed that the prediction model learned pattern transition from image features, even though the continuous and cyclical nature of the stages, which is a characteristic of spermatogenesis, was not explicitly provided as a teacher signal or as teacher knowledge (Figure 5). A mechanism that can actively utilize this feature of the continuous and cyclical nature of the seminiferous tubule stages in learning can be devised to further improve prediction accuracy.

In recent years, deep learning methods have been proposed for use in infertility treatment for the evaluation of sperm, oocytes, and fertilized embryos[23, 24, 25, 26]. Although our method is likely to be used mainly in research into understanding spermatogenesis, it may also play a role in clinical practice by offering a more detailed or precise way to evaluate sperm quality and production, potentially leading to improved diagnosis and treatment of male infertility. In addition, our method could also be applied to medical research where reproductive toxicity is less pronounced and overlooked, such as the effects of aging, diet and environmental factors on spermatogenesis.

## Data availability

Part of the data used in this paper is available at https://github.com/funalab/STSP. The rest of the data will be handled by upon request.

## Code availability

The code used in this paper is available at https://github.com/funalab/STSP.

## Acknowledgements

The authors thank the suggestions and discussions with Takahiro G. Yamada in Funahashi-lab, Keio University and Daisuke Mashiko in Research Institute for Microbial Diseases, Osaka University. This research was funded by a JST CREST grant to M.I. and A.F. (Grant Number JPMJCR21N1). The NVIDIA Tesla V100 was used in the miniRAIDEN computer server owned by RIKEN.

## Author contributions

Y.T., M.I., and A.F. designed the conceptual idea and the study. Y.T. and T.M. implemented the classification algorithm. T.E., Y.H., and M.I. provided the datasets of seminiferous tubules. Y.T., T.E., T.M., M.I., and A.F. wrote the manuscript, with suggestions from the other authors. Y.T. and T.E. contributed equally to this work. A.F. and M.I are the co-corresponding authors of this study.

