## Supplementary Material for "Deep learning–based automated prediction of mouse seminiferous tubule stage by using bright-field microscopy"

### Supplementary Figures

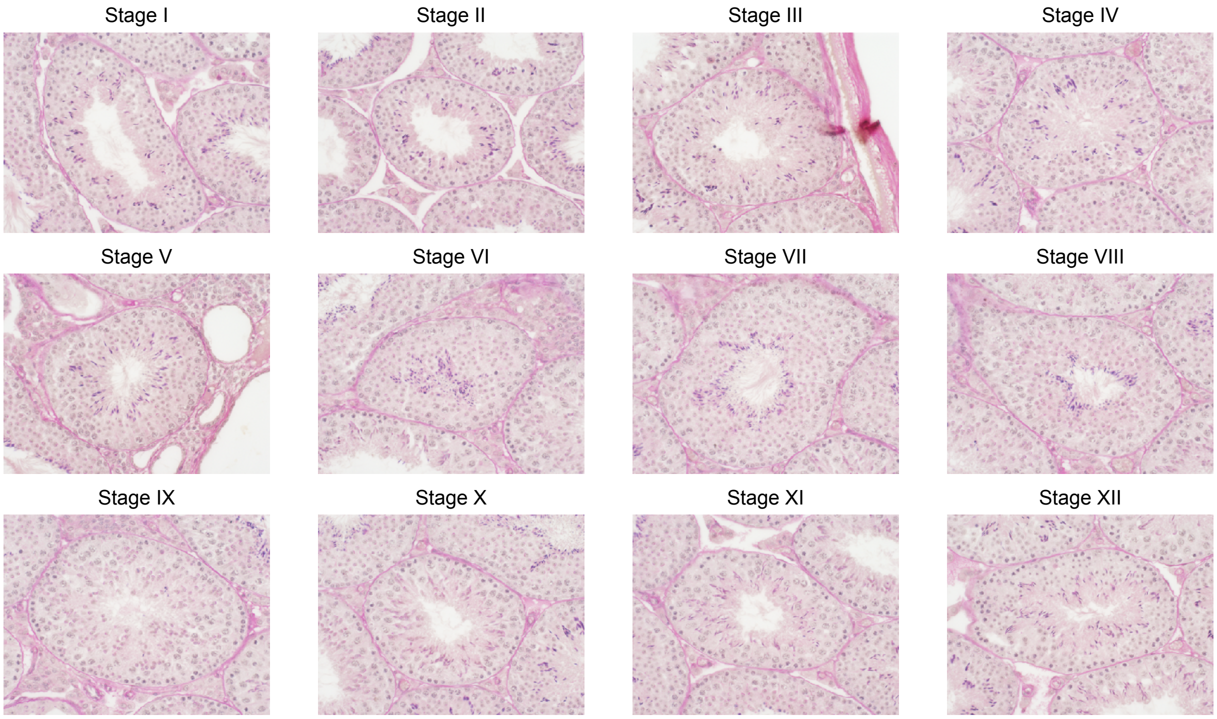


Supplementary Figure S1: Typical fine seminiferous tubules tissue images, by stage. Typical seminiferous tubule tissue images from Stages I to XII are shown. The seminiferous tubule stage associated with an image is assigned with respect to the tubule in the center of the image.

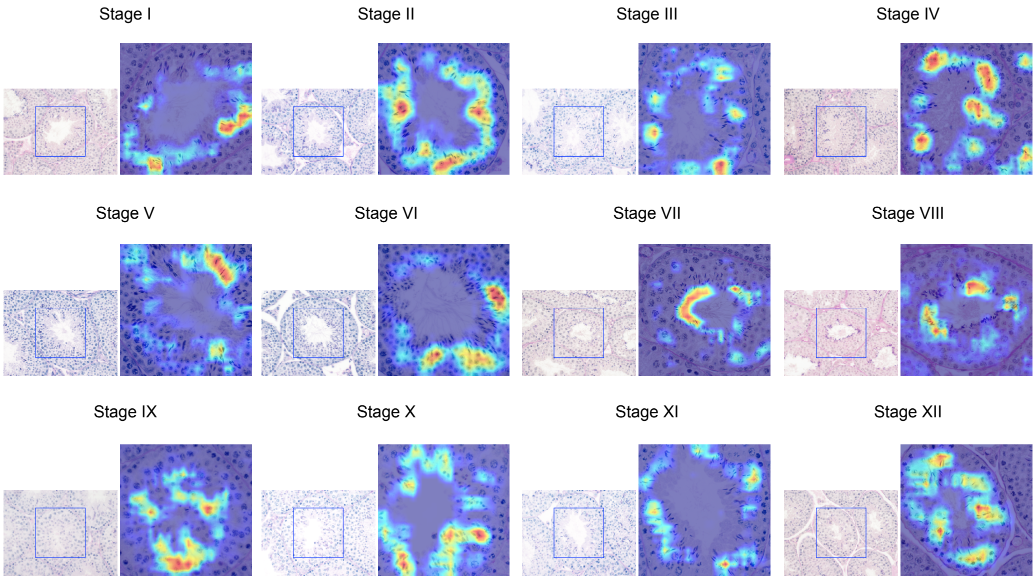


Supplementary Figure S2: Image features in each stage that form the basis for seminiferous tubule stage prediction.

Representative features of the seminiferous tubule images that were used as the basis for stage prediction by ResNet-50 model trained on the seminiferous tubules dataset, visualized by Grad-CAM. All of these results represent the image features that were used as the basis for correct stage prediction. Higher activation in the heat map (indicated by warmer colors) indicates a stronger contribution to the prediction. Stage I focused on step 1 spermatids (round spermatids) and step 13 spermatids (elongated spermatids). The "step" means a seminiferous epithelium cycle. Stage Ⅱ focused on step 2 spermatids (round spermatids) and step 14 spermatids (elongated spermatids). Stage Ⅲ focused on step 3 spermatids (round spermatids) and step 14 spermatids (elongated spermatids). Stage Ⅳ focused on step 4 spermatids (round spermatids), step 15 spermatids (elongated spermatids), and pachytene spermatocytes. Stage V focused on step 5 spermatids (round spermatids) and step 15 spermatids (elongated spermatids). Stage Ⅵ focused on step 6 spermatids (round spermatids) and step 15 spermatids (elongated spermatids). Stage Ⅶ focused on step 7 spermatids (round spermatids), step 16 spermatids (elongated spermatids), and pachytene spermatocytes. Stage Ⅷ focused on step 8 spermatids (round spermatids) and step 16 spermatids (elongated spermatids). Stage Ⅸ focused on step 9 spermatids (elongated spermatids). Stage X focused on step 10 spermatids (elongated spermatids) and pachytene spermatocytes. Stage Ⅺ focused on step 11 spermatids (elongated spermatids) and diplotene spermatocytes. Stage Ⅻ focused on step 12 spermatids (elongating spermatids), meiotically dividing spermatocytes, and late zygotene spermatids.

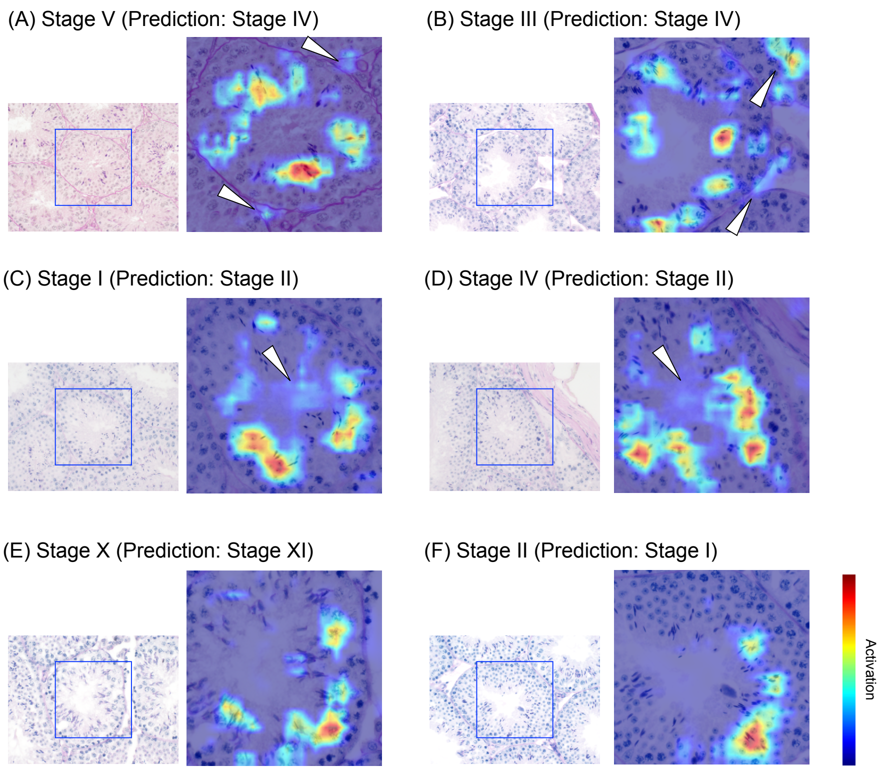


Supplementary Figure S3: Image features used as the basis for failed predictions of seminiferous tubule stage.
Representative features of the seminiferous tubule images that were used as the basis for stage prediction by ResNet-50 model trained on the seminiferous tubules dataset, visualized by Grad-CAM. All of these results represent image features that failed to correctly predict the stage. Higher activation in the heat map (indicated by warmer color) indicates a stronger contribution as a basis for prediction. (A) Features that were the focus of attention when a prediction of Stage IV was made for a Stage V image. As indicated by the arrowheads, attention was paid to image features other than the target (centered) seminiferous tubules, which were the target of the prediction. (B) Features focused on when a prediction of Stage IV was made for a Stage III image. As indicated by the arrowheads, the features focused on were the same as in (A). (C) Features that were focused on when a prediction of Stage II was made for a Stage I image. As indicated by the arrowheads, attention was paid to the lumen of the seminiferous tubules where spermatogonia, spermatocytes, and spermatids are not present. (D) Features that were focused on when a prediction of Stage II was made for a Stage IV image. The features focused on were the same as in (C). (E) Features that were focused on when a prediction of Stage XI was made for a Stage X image. Only some of the specific cells surrounding the lumen of the seminiferous tubules were focused on. (F) Features that were focused on when a prediction of Stage I was made for a Stage II image. As in (E), only a portion of the specific cells surrounding the lumen of the seminiferous tubules were focused on.

### Supplementary Tables

Supplementary Table S1: Hyperparameters used to train each neural network.

| Hyperparameter | Value |
|---|---|
| Number of epochs | 5,000 |
| Adam learning rate | 0.001 |
| Adam betas | (0.9, 0.999) |
| Adam eps | $1.0 \times 10^{-8}$ |
| Mini-batch size | 12 |
| Weight decay | 0.0005 |

Supplementary Table S2: Comparison of the Balanced Accuracy of each model in predicting seminiferous tubule stage on the basis of four-fold cross-validation. Bold type indicates the maximum value at each fold and mean.

| Model | fold1 | fold2 | fold3 | fold4 | Mean (S.D.) |
|---|---|---|---|---|---|
| ResNet-50 | 0.8111 | 0.7942 | **0.7994** | 0.7546 | 0.7898 (0.0212) |
| ResNeXt-50 | **0.8397** | **0.8399** | 0.7992 | 0.7907 | **0.8174** (0.0226) |
| WideResNet-50 | 0.8161 | 0.7897 | 0.7819 | **0.7990** | 0.7967 (0.0127) |
| MobileNet-v3 | 0.8040 | 0.7935 | 0.7839 | 0.7649 | 0.7866 (0.0144) |
